# Neural Activation of Human Primary Intestinal Enteroendocrine L-cells

**DOI:** 10.64898/2026.11.03.763680

**Authors:** Jack Lindsay, Daniel Zeve, Sara Rudolph, Amy Shepherd, Allison Grossberg, David Breault, Michael Levin, Donald Ingber

**Author notes:** Address correspondence to: Donald Ingber, MD,PhD (em.

## Abstract

Intestinal enteroendocrine L-cells play a central role in metabolic homeostasis by secreting the gut hormones GLP-1 and PYY, but whether and how neural signals regulate human L-cell function remains poorly understood. This gap reflects the limited availability of human model systems enriched for mature L-cells. Here, we establish a differentiation strategy that generates human donor-derived ileal organoids with markedly increased L-cell abundance, enabling functional interrogation of L-cell physiology in their basal state. Using this system, we show that cholinergic and adrenergic stimuli activate L-cells, triggering calcium influx and secretion of GLP-1 and PYY. Pharmacological analysis reveals that cholinergic regulation of human L-cells is mediated predominantly through M3 muscarinic receptors, while adrenergic regulation is mediated through the β1 adrenergic receptor.

Neurotransmitter-evoked activation also initiates gap junction–dependent calcium waves that propagate across the intestinal epithelium; however, disruption of epithelial coupling by abolishing calcium propagation does not impair hormone secretion. Together, these findings identify L-cells as integrative hubs that couple neural signals to gut hormone secretion and epithelial calcium waves, providing mechanistic insight into the gut-brain axis and how the nervous system regulates intestinal endocrine function in human intestine.

## Introduction

The intestinal epithelium is a major endocrine organ that plays a central role in metabolic homeostasis through the secretion of gut hormones that regulate nutrient absorption, insulin release, appetite, and energy balance^1–4^. Among these hormones, glucagon-like peptide-1 (GLP-1) and peptide YY (PYY) have emerged as key regulators of glucose metabolism and satiety, and they form the basis of widely used therapies for diabetes and obesity^5–10^. In the intestine, enteroendocrine L-cells are the principal source of GLP-1 and PYY^11–13^, positioning them as critical control points in the gut–brain axis that links nutrient availability to systemic metabolic responses^14–16^.

L-cells are classically viewed as nutrient sensors that respond directly to luminal glucose, amino acids, and fatty acids^17–20^. Following nutrient stimulation, L-cells release peptide hormones that can be directly sensed by both intrinsic and extrinsic nerve fibers that are apposed to the basal surface of the intestinal epithelium^21,22^. However, L cells also express receptors for a variety of neurotransmitters and neuromodulators^23^, opening the possibility that this communication may be bidirectional. These observations suggest that the nervous system may act as a mediator of L-cell function, but thus far direct functional evidence for neural regulation of L-cells in humans remains limited.

Existing mechanistic insights into neurotransmitter control of L-cell function derive from animal studies or immortalized neuroendocrine cell lines. These studies suggest that cholinergic and adrenergic signaling can stimulate GLP-1 and PYY release through muscarinic, nicotinic, and adrenergic receptors^24–29^. However, whether these pathways operate in human L-cells and how they drive downstream signaling across the intestinal epithelium are unresolved questions. Progress in addressing these issues has been constrained by the scarcity of mature L-cells in existing human intestinal organoid models, which could enable direct functional assays.

Beyond local neural inputs, L-cells may also respond to systemic signals that reflect organismal metabolic state. Circulating catecholamines, such as epinephrine, rise rapidly during stress and arousal and exert broad effects on glucose metabolism and energy mobilization^30,31^. While epinephrine is best known for its endocrine actions, the expression of adrenergic receptors on L-cells raises the possibility that these cells act as sensors of the autonomic state, coupling systemic neuroendocrine signals to intestinal hormone secretion. Whether and how such signals directly engage human L-cells has not been established.

Here, we uncover neural pathways that activate primary human L-cells by first establishing a differentiation strategy that markedly increases L-cell abundance in human donor-derived ileal organoids to produce a tractable model to characterize hormone secretion in response to neurotransmitters. Then we use this system to identify cholinergic and adrenergic pathways that drive calcium responses, promote GLP-1 and PYY secretion, and initiate gap junction–dependent calcium waves that propagate across the intestinal epithelium. Our findings position L-cells as integrative hubs that translate autonomic and neuroendocrine inputs into coordinated epithelial and endocrine responses, providing mechanistic insight into how neural signaling is coupled to gut hormone secretion in humans.

## Results

To establish a human ileal organoid model that enables functional studies of L-cells, we adapted the previously described G2D12 differentiation regimen in which the organoids are cultured for 2 days in growth medium followed by 12 days in base differentiation medium. G2D12 was previously shown to induce chromogranin A (CHGA)-positive enteroendocrine cells as well as support the production and secretion of GLP-1 and PYY in human rectal organoids^32^. To optimize L-cell differentiation in human donor-derived ileal organoids, we supplemented G2D12 with 1 μM EPZ-6438 (EPZ; also known as tazemetostat), a selective inhibitor of EZH2, the catalytic subunit of Polycomb repressive complex 2 (PRC2) ^33–35^. EZH2 was explored as a potential induce of L-cell differentiation because PRC2 inhibition has been shown to promote secretory differentiation and patients on EPZ-6438 reported side effects similar to those in patients receiving GLP-1R agonists^33,35,36^.

Relative to G2D12 alone, EPZ supplementation increased expression of the L-cell hormone genes GCG and PYY by approximately 2-fold (**Fig. 1A**). Following stimulation with 10 μM forskolin and 10 μM IBMX to increase cAMP and mobilize stores of secretory vesicles, the EPZ-differentiated organoids secreted significantly more total GLP-1 and PYY than plate-matched G2D12 cultures across three ileal organoid lines (**Fig. 1B**). Consistent with these transcriptional and secretory measurements, immunofluorescence microscopic analysis revealed a marked increase in CHGA-positive and GLP-1-positive cells in EPZ-treated organoids (**Fig. 1C**). Quantification of L-cells relative to total nuclei confirmed an increase in L-cell abundance in each matched comparison, with L-cells comprising roughly up to 3% of cells in EPZ-differentiated organoids compared with no more than approximately 1% in G2D12 controls (**Fig. 1D**). Thus, EPZ supplementation increased both the abundance of ileal L-cells and the total hormone output in these human intestinal organoid cultures.

**Fig. 1.**
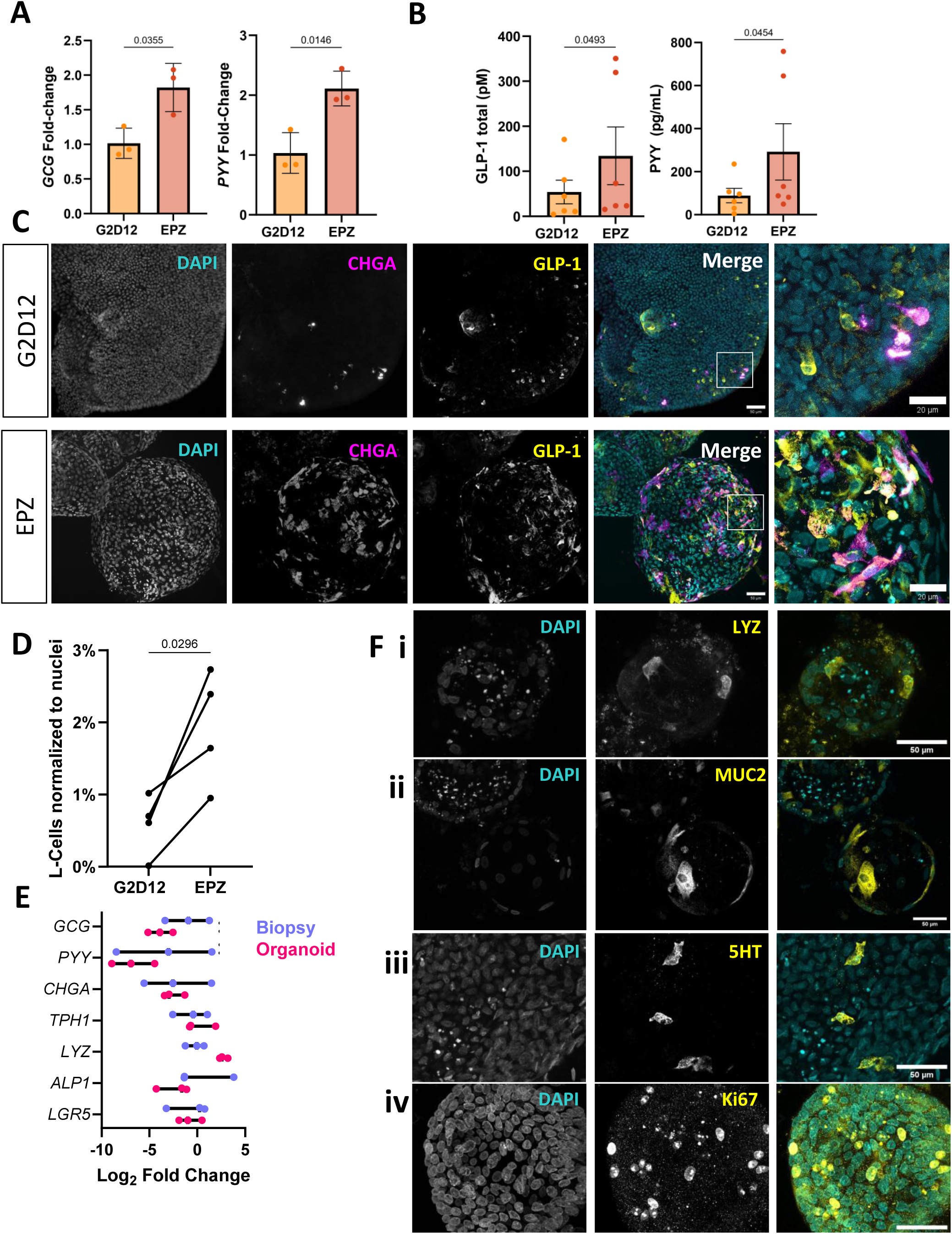
Generation of human ileal organoids enriched for functional L-cells. **(A)** Expression of marker genes for L-cells proglucagon (GCG) and PYY in ileal organoids cultured under G2D12 and EPZ conditions (Lines indicate SD, comparison by Welch’s t-test, n = 3, one organoid line). **(B)** Secretion of GLP-1 and PYY proteins during incubation with 10 μM forskolin and 10 μM IBMX in ileal organoids cultured under G2D12 and EPZ conditions. Each dot represents the average of 4 independent wells on a single plate, statistics performed on matched groups of wells run on the same plate (Lines indicate SEM, comparison by ratio paired t-test, n = 6 across 3 organoid lines). **(C)** Representative images of organoids differentiated in either G2D12 or EPZ stained for nuclei (DAPI), EEC marker CHGA, and L-cell marker GLP-1. Scale bar = 50 µm. **(D)** Relative number of L-cells in organoids differentiated in either G2D12 or EPZ identified via immunocytochemistry. Lines connect matches pairs of organoid wells grown on the same plate. Each data point represents images of >5000 nuclei. (comparison by paired t-test, n = 4 across 3 organoid lines. Scale bar = 50 µm). **(E)** Relative expression of intestinal epithelial markers between three healthy patient biopsies and organoids differentiated in EPZ. All points compared to average biopsy expression. (n = 3, organoids from 2 lines) **(F)** Immunocytochemistry of cell types present in organoids differentiated with EPZ: (i) Paneth cells marked by LYZ, (ii) goblet cells marked by MUC2, (iii) enterochromaffin cell marked by 5HT, (iv) proliferating cells marked by Ki67. Scale bar = 50 microns.

Because PRC2 inhibition could broadly alter epithelial differentiation, we next assessed whether EPZ-differentiated organoids retained representation of the major epithelial lineages of the ileum. EPZ organoids expressed markers of L-cells (GCG and PYY), the broader enteroendocrine lineage (CHGA), enterochromaffin cells (Tryptophan hydroxylase 1, TPH1), Paneth cells (Lysozyme, LYZ), absorptive enterocytes (intestinal alkaline phosphatase, ALPI), and intestinal stem or progenitor cells (Leucine Rich Repeat Containing G Protein-Coupled Receptor 5, LGR5) when benchmarked against primary ileal biopsies using qPCR (**Fig. 1E**). The range of gene expression in EPZ-differentiated organoids largely overlapped with that observed in clinical biopsies, with the exception of LYZ which appeared to be overexpressed in organoids (**Fig. 1E**). Immunofluorescence staining confirmed the presence of LYZ-positive Paneth cells, MUC2-positive goblet cells, 5HT-positive enterochromaffin cells, and Ki67-positive proliferating cells within EPZ-differentiated organoids (**Fig. 1F**), representing all the major epithelial populations found in the human ileum^37^. Together, these results establish that EPZ supplementation enriches functional L-cells while still supporting multilineage epithelial differentiation, and thus providing a tractable human ileal organoid model for mechanistic studies of L-cell signaling and secretion.

We used this L-cell enriched human ileal organoid model to evaluate whether candidate neural mediators alter intestinal L-cell activity. Transcriptomic profiling in sorted human L-cells has identified a broad repertoire of G protein-coupled receptors and ion channels, while intracellular Ca²⁺ imaging has been shown to be a sensitive readout of L-cell excitation in cultured murine L-cells^18,23,38^. We therefore performed live Ca²⁺ imaging in EPZ-differentiated ileal organoids, polarized to expose their basal membranes to neurotransmitters (**Fig. 2A)**. When multiple candidate mediators including acetylcholine, epinephrine, norepinephrine, GABA, and VIP were added to the organoids, only acetylcholine and epinephrine significantly increased whole-organoid Ca²⁺-dependent fluorescence relative to vehicle (**Fig. 2B**). Norepinephrine appeared to produce a small response but it did not reach statistical significance, whereas GABA and VIP did not elicit any detectable responses. Because acetylcholine and epinephrine were the only ligands to evoke robust Ca²⁺ responses, we focused subsequent analyses on these two neurotransmitters.

**Fig. 2.**
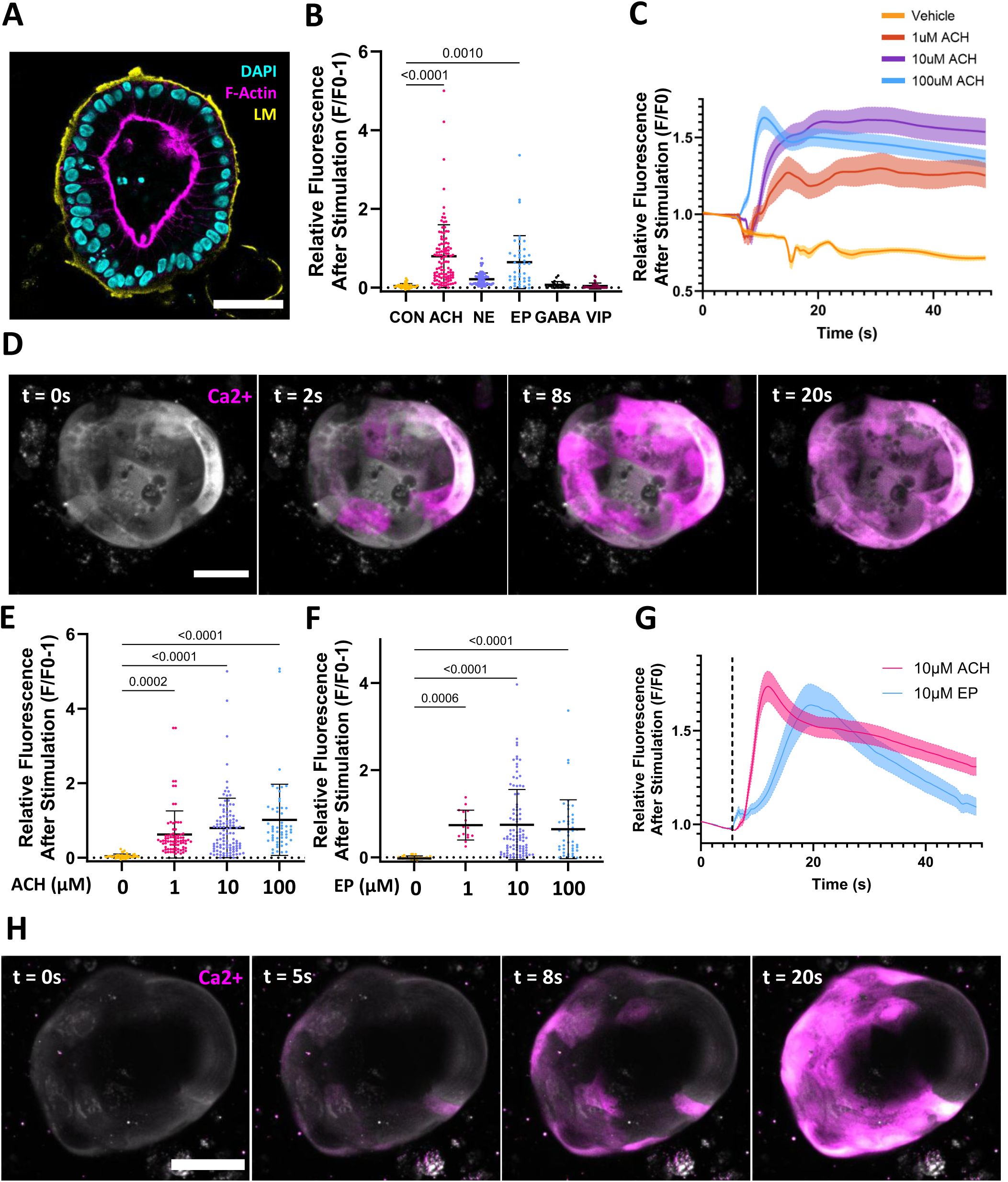
Cholinergic and adrenergic signals evoke a calcium wave in ileal organoids. **(A)** Representative confocal image of an ileal organoid after calcium imaging showing apical-in polarization. LM: Laminin **(B)** Peak change in fluorescence from baseline after stimulation with vehicle (CON), norepinephrine (NE), epinephrine (EP), Gamma-Aminobutyric Acid (GABA), vasoactive intestinal peptide (VIP) or acetylcholine (ACH) (Lines indicate SD, comparison by one-way ANOVA with Dunnett’s multiple comparison test versus CON, nonsignificant results not shown, n = 23 – 78, 3-5 independent wells per condition). **(C)** Average traces for changes in calcium levels after stimulation with either vehicle or escalating doses of acetylcholine. Shaded areas represent standard error of the mean. **(D)** Representative maximum intensity projection image of a single organoid after stimulation with 10µm acetylcholine. Greyscale image is calcium at baseline, pink is changes in calcium from baseline. Scale bar = 25 µm. **(E)** Effect of acetylcholine dose on peak change in fluorescence from baseline (Lines indicate SD, comparison by one-way ANOVA with Dunnett’s multiple comparison test versus vehicle, nonsignificant results not shown, n = 45 – 107, 3-4 independent wells per condition across 2 organoid lines). **(F)** Effect of epinephrine dose on peak change in fluorescence from baseline (Lines indicate SD, comparison by one-way ANOVA with Dunnett’s multiple comparison test versus vehicle, nonsignificant results not shown, n = 15 – 97, 4 independent wells per condition across 3 organoid lines) **(G)** Comparison of the time response to stimulation with either 10μM acetylcholine or 10μM epinephrine, shaded areas represent standard error of the mean, n= 37 – 97 **(H)** Representative maximum intensity projection image of a single organoid after stimulation with 10 µm epinephrine. Greyscale image is calcium at baseline, pink is changes in calcium from baseline. Scale bar = 25 µm.

Acetylcholine significantly increased Ca²⁺-dependent fluorescence in a dose-dependent manner when its concentration was raised from 1 and 100 μM (**Fig. 2C,E**). At 10 μM, the response began within seconds of ligand addition, reached an early peak, and remained elevated above baseline throughout the post-stimulation imaging period (**Fig. 2C**). Time-lapse imaging showed that acetylcholine-evoked fluorescence initially appeared in cells within spatially restricted regions and then rapidly expanded across the organoid epithelium, producing a wave-like response (**Fig. 2D**).

Epinephrine likewise elicited significant Ca²⁺ responses across the 1–100 μM concentration range, although the response magnitude did not increase monotonically with concentration (**Fig. 2F**). At 10 μM, the mean epinephrine response rose more gradually and reached its peak later than the response to acetylcholine, followed by a more pronounced decline toward baseline during the remainder of the imaging period (**Fig. 2G**). Time-lapse imaging revealed a similar spatially expanding pattern, in which epinephrine-responsive regions emerged locally before involving a broader portion of the epithelium, but with slower apparent kinetics than the acetylcholine response (**Fig. 2H**).

Together, these results identify acetylcholine and epinephrine as acute triggers of robust but kinetically distinct epithelial Ca²⁺ responses in human ileal organoids. Because these responses encompassed substantially larger regions of the epithelium than would be predicted from the L-cell abundance we measured (**Fig. 1**), their spatial dynamics suggested that a rare sensor population may initiate a signal that subsequently propagates to neighboring epithelial cells.

Having identified acetylcholine as a potent stimulus of epithelial Ca²⁺ signaling, we next asked whether this response was mediated by muscarinic or nicotinic acetylcholine receptors. Previous studies in rat intestinal preparations and the human NCI-H716 enteroendocrine cell line demonstrated that muscarinic receptor activation can stimulate GLP-1 secretion, although the receptor subtypes implicated differed between the experimental models^24,25^. The nonselective muscarinic agonist bethanechol reproduced the Ca²⁺ response elicited by acetylcholine in our ileal organoids, whereas pretreatment with the nonselective muscarinic antagonist atropine markedly reduced the response to acetylcholine (**Fig. 3A,B**). In contrast, the stimulation of the nicotinic receptor by addition of nicotine did not produce a detectable Ca²⁺ response under the conditions tested (**Fig. 3A, B**). These results indicate that the acute response to acetylcholine is mediated predominantly by muscarinic rather than nicotinic receptor signaling.

**Fig. 3.**
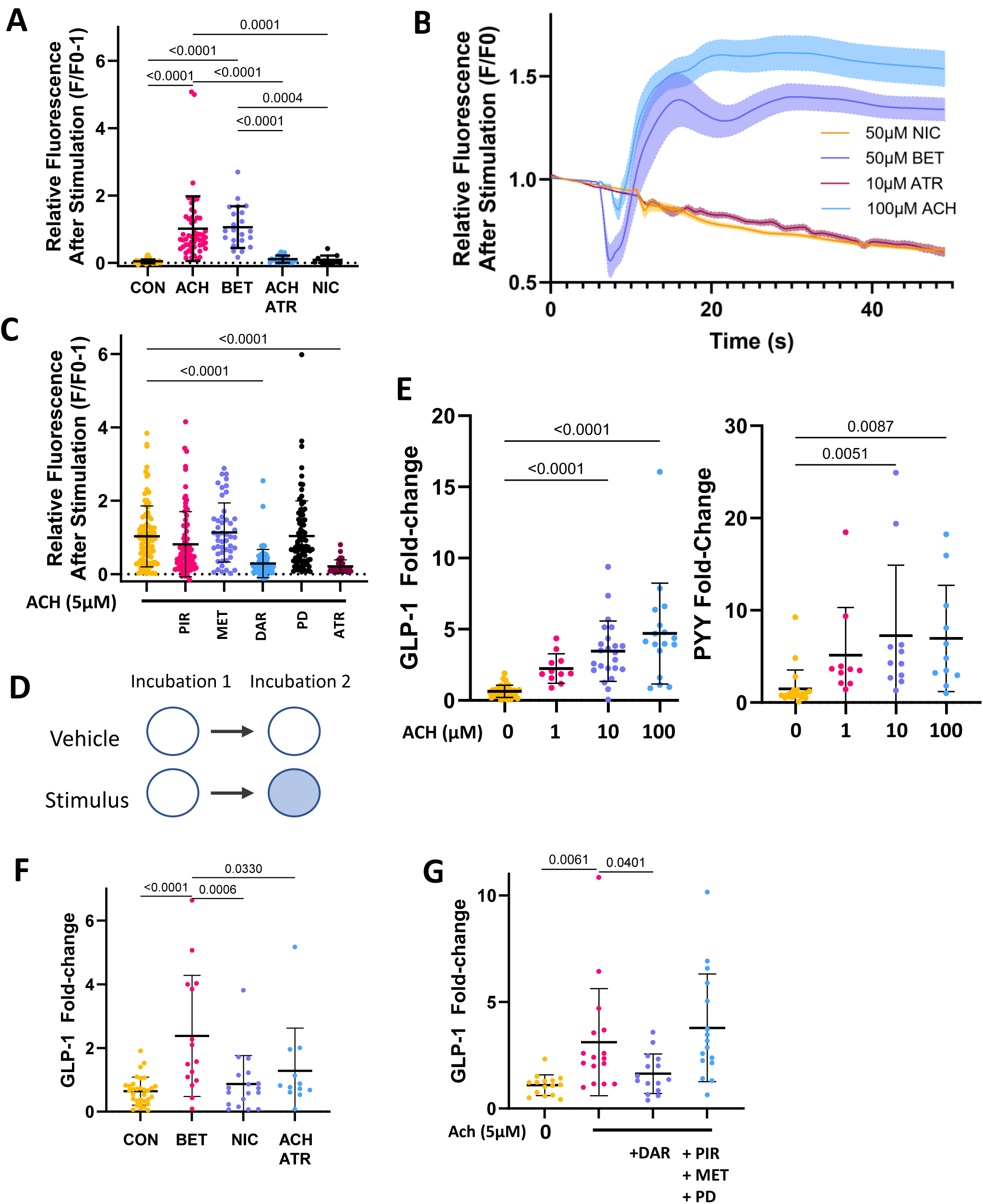
Muscarinic signaling couples cholinergic input to GLP-1 and PYY secretion in human L-cells. **(A)** Peak change in calcium levels following stimulation with either 100μM acetylcholine (ACH), muscarinic receptor agonist 50μM bethanechol (BET), 100μM ACh during incubation with muscarinic antagonist 10 μM atropine (ATR), or nicotinic receptor agonist 50 μM nicotine (NIC) (Lines indicate SD, comparison by one-way ANOVA with Dunnett’s multiple comparison test versus CON, nonsignificant results not shown, n = 11 – 54, three independent wells per condition from one line). **(B)** Average calcium traces after treatment with nicotine (NIC), bethanechol (BET), 100 μM acetylcholine during incubation with atropine (ATR), or acetylcholine (ACH). Dotted lines represent SEM, n = 8 – 22, one line. **(C)** Peak change in calcium from baseline after stimulation with acetylcholine (ACh) during incubation with either M1 antagonist pirenzepine, 100 nM (PIR), M2 antagonist methoctramine, 100nM (MET), M3 antagonist darifenacin, 10 nM (DAR), M4 antagonist PD 102807, 300nM (PD), or non-selective muscarinic antagonist atropine, 10 μM (ATR) (Lines indicate SD, comparison by one-way ANOVA with Dunnett’s multiple comparison test versus 5μM ACh, nonsignificant results not shown, n = 66 – 94 from 4 independent wells per condition across two organoid lines). **(D)** Incubation scheme for ELISA. **(E)** Effects of acetylcholine on GLP-1 and PYY secretion. (Lines indicate SD, comparison by one-way ANOVA with Dunnett’s multiple comparison test versus vehicle, nonsignificant results not shown, n = 9 – 22 across two organoid lines)**. (F)** Effects of bethanechol, 100 μM (BET), nicotine, 100μM (NIC), and 10 μM acetylcholine (ACh) during incubation with atropine, 10 μM (ATR) on GLP-1 secretion. (Lines indicate SD, comparison by one-way ANOVA with Dunnett’s multiple comparison test versus BET, n = 12 – 29 across two organoid lines). **(G)** Effects of muscarinic antagonists on GLP-1 secretion. (Lines indicate SD, comparison by multiple Welch’s t-tests, nonsignificant results not shown, n = 15 across two organoid lines.)

We next used subtype-preferring muscarinic antagonists to determine which receptor subtype contributed to the acetylcholine response. Organoids were pretreated with pirenzepine, methoctramine, darifenacin, or PD 102807 before stimulation with acetylcholine to identify which muscarinic receptors mediate its responses. An acetylcholine dose of 5 μM was used because it consistently produced a robust response without overwhelming the effects of the antagonists at their selective concentrations. Darifenacin, an M3 muscarinic receptor-preferring antagonist, significantly attenuated the acetylcholine-evoked Ca²⁺ response, producing an effect similar to that of atropine (**Fig. 3C**). By contrast, pretreatment with pirenzepine, methoctramine, and PD 102807, which respectively target M1, M2, and M4 muscarinic receptors, did not significantly reduce the response. This pharmacological profile implicates signaling through the M3 receptor as mediating acetylcholine-evoked epithelial Ca²⁺ activity in human L-cells.

We next determined whether acetylcholine-induced Ca²⁺ signaling was accompanied by secretion of L-cell hormones using a sequential, within-well, secretion assay (**Fig. 3D**). Importantly, addition of acetylcholine stimulated secretion of both GLP-1 and PYY, with statistically significant increases observed at 10 and 100 μM relative to the control condition (**Fig. 3E**). Bethanechol also stimulated GLP-1 secretion, whereas nicotine did not, consistent with the M3 receptor-class dependence observed by Ca²⁺ imaging (**Fig. 3F**). Finally, darifenacin, a competitive antagonist with strong preference for M3, significantly reduced acetylcholine-evoked GLP-1 secretion, whereas combined inhibition of M1-, M2-, and M4 receptors with competitive antagonists did not reproduce the inhibitory effect of darifenacin (**Fig. 3G**). Together, the Ca²⁺ imaging and hormone secretion experiments independently demonstrate that acetylcholine activates a muscarinic signaling pathway within human ileal organoids and provide convergent pharmacological evidence favoring a predominant role for the M3 receptor in coupling cholinergic stimulation to GLP-1 release in human L-cells.

We next sought to understand whether the calcium dynamics observed after stimulation with epinephrine triggered GLP-1 and PYY secretion, and to identify the receptor subtypes responsible. Epinephrine increased secretion of both GLP-1 and PYY, with significant responses observed at 10 μM for GLP-1 and both 1 and 10 μM for PYY (**Fig. 4A**). These findings confirmed that the Ca²⁺ response to epinephrine is accompanied by increased secretion of both major L-cell hormones. To identify potential adrenergic receptors that mediate this response, we analyzed receptor expression across four publicly available transcriptomic datasets containing primary human tissue- or organoid-derived enteroendocrine cells^23,39–41^. Four GEO datasets were analyzed to profile adrenergic receptor expression in human GCG+ L-cells (**Fig. 4B)**. GSE284419 (single-cell, ileal organoid, CHGA-sorted) provides primary expression profiling of L-cells against other EECs. GSE148224 (bulk, ileal organoid) and GSE114853 (bulk, primary jejunum and ileum) provide differential expression against the surrounding epithelium, with GSE114853 additionally supporting the within-EEC comparison. GSE146799 (bulk CEL-Seq2, organoid) provides sorted reporter populations and contributes both comparisons.

**Fig. 4.**
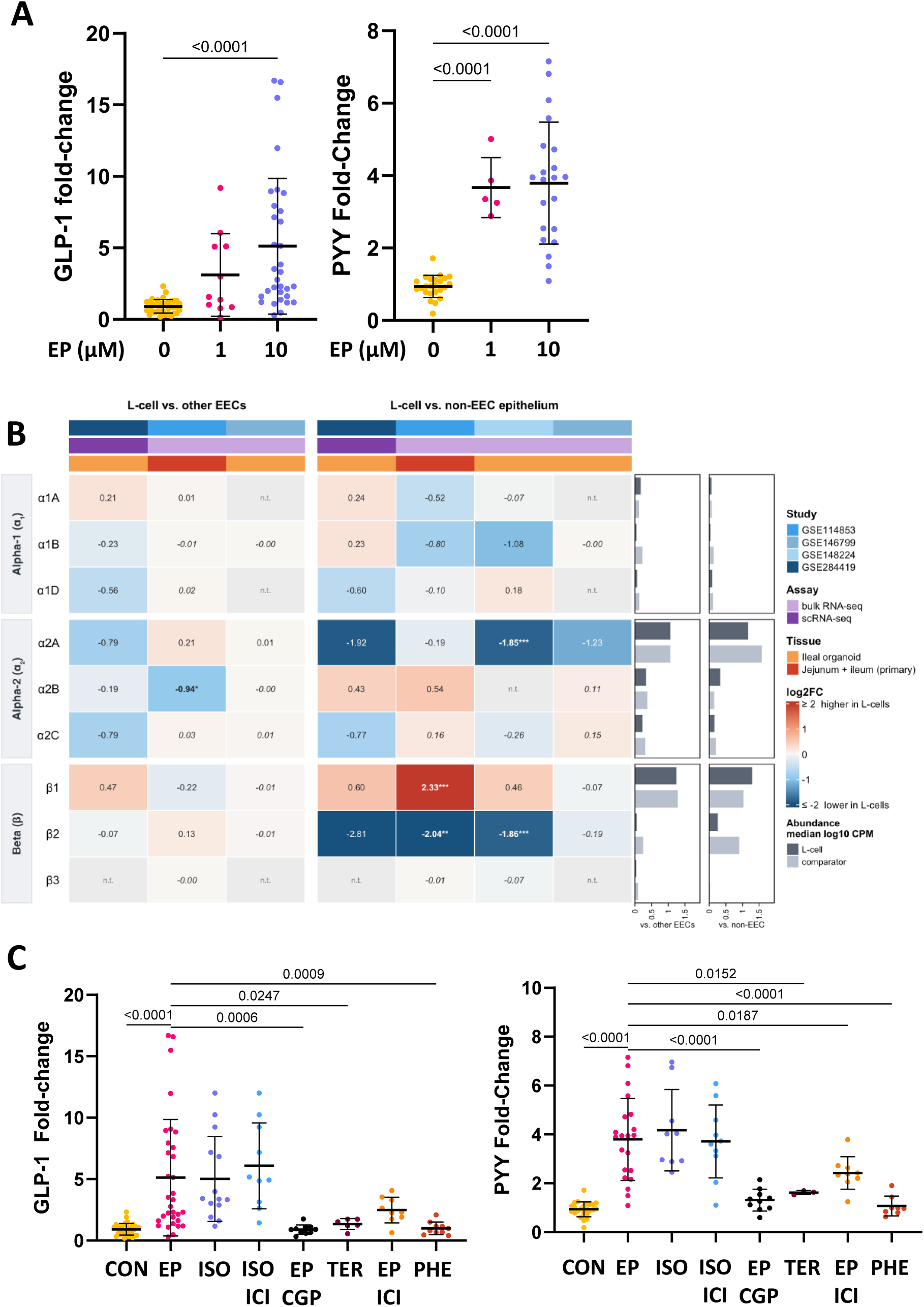
β1-adrenergic signaling promotes GLP-1 and PYY secretion from human ileal organoids. **(A)** Within-well fold change in GLP-1 and PYY secretion following stimulation with the indicated concentrations of epinephrine (EP) (Lines indicate SD, comparison by one-way ANOVA with Dunnett’s multiple-comparison test versus vehicle; nonsignificant results not shown, n = 4–37 across three donor-derived organoid lines). **(B)** Adrenergic receptor log2 fold changes in human L-cells across four GEO datasets. Rows, nine adrenergic receptors grouped by subfamily; columns, one color per dataset, split into L-cells versus other EECs (left) and versus non-EEC epithelium (right). Red, higher in L-cells; blue, lower; color represents ±2 log2 units, with exact values printed in each cell. Values are shrunken fold-change estimates. Asterisks represent Benjamini-Hochberg adjusted p-values: *p < 0.05, **p < 0.01, ***p < 0.001. Values in italic had no adjusted p-value after DESeq2 independent filtering and are therefore untested. Genes absent from a particular dataset are shown as “n.t.”. p-values are not shown for GSE284419 as this experiment included only a single donor. Right bar plot panels show the abundance of each gene as median log10(CPM + 1) for L-cells (dark) and the block’s comparator (light), computed only from datasets containing both groups. **(C)** Effects of adrenergic agonists and antagonists on GLP-1 and PYY secretion. Vehicle (CON), epinephrine, 10 μM (EP), β1/β2-adrenergic receptor agonist isoproterenol, 1 μM (ISO), β1/β2-adrenergic receptor agonist isoproterenol during incubation with β2-selective antagonist antagonist ICI 118,551, 100 nM (ICI), epinephrine during incubation with β1-selective antagonist CGP 20712A, 100 nM (CGP), β2-adrenergic receptor agonist terbutaline, 10μM (TER), epinephrine during incubation with ICI 118,551, α2-adrenergic receptor agonist phenylephrine, 10 μM (PHE) (Lines indicate SD, comparison by one-way ANOVA with Dunnett’s multiple comparison test versus epinephrine, nonsignificant results not shown, n = 6 – 37 across three organoid lines).

The most consistent finding across these datasets is that the adrenergic α2A receptor (ADRA2A) is depleted in L-cells. Against non-EEC epithelium it is strongly depleted in GSE148224 (z = -7.00, p = 4.3 × 10−11, log2FC = -1.85 ± 0.27; mean CPM = 18.2 vs 56.0, n = 8 vs 8) and negative in all four datasets, though not significantly so in primary tissue (z = -0.59, p = 0.688, log2FC = -0.19 ± 0.39). In the single cell data, it is detected in 8.8% of L-cells compared to 18.0% of other EECs and 34.4% of non-EEC cells.

Likewise, the β2 receptor (ADRB2) is significantly depleted in L-cells relative to non-EEC epithelium in both bulk datasets (GSE114853, z = -3.17, p = 0.005, log2FC = -2.04 ± 0.77; GSE148224, z = -4.27, p = 1.1 × 10−4, log2FC = -1.86 ± 0.49) and is flat against other EECs (z = 1.63, p = 0.369, log2FC = 0.13 ± 0.83). In the single cell data, it is detected in 0.2% of L-cells against 15.3% of non-EEC cells.

Importantly, this analysis revealed that the adrenergic β1 receptor (ADRB1) is significantly enriched in L-cells relative to non-EEC epithelium but not relative to other EECs. In primary tissue, it is enriched over epithelium (z = 4.88, p = 7.1 × 10−6, log2FC = 2.33 ± 0.52, mean CPM = 24.2 vs 3.1, n = 13 vs 13) yet flat against other EECs in the same dataset and model (z = -1.61, p = 0.379, log2FC = -0.22 ± 0.50). Two of the three organoid datasets showed the same pattern without reaching significance (GSE148224, z = 1.62, p = 0.200, log2FC = 0.46 ± 0.33). In contrast, the β3 receptor (ADRB3) was essentially undetectable, at or below 0.07 CPM in L-cells in every dataset. The α1A, α1B, α1D and α2C receptors were likewise expressed below 0.5 CPM in L-cells and show no significant differential expression in any contrast. α2B is the one exception among the low-abundance receptors: it was significantly depleted against other EECs in primary tissue (z = -3.12, p = 0.022, log2FC = -0.94 ± 0.70).

Given the relative abundance and enrichment of ADRB1 and ADRB2, we pharmacologically manipulated GLP-1 and PYY secretion using chemical agonists and antagonists against these receptors. As expected, the nonselective β-adrenergic agonist isoproterenol stimulated GLP-1 and PYY secretion to levels comparable to epinephrine, supporting a β-adrenergic mechanism (**Fig. 4C**). In contrast, the β2 receptor-preferring agonist terbutaline and the α1 adrenergic receptor agonist phenylephrine produced substantially smaller responses. Interestingly, the β1-selective antagonist CGP 20712A markedly reduced epinephrine-stimulated GLP-1 and PYY secretion, whereas selective antagonism of the β2 receptor by ICI 118,551 produced a more limited effect. Consistent with this pattern, isoproterenol remained stimulatory in the presence of ICI 118,551. Together, the transcriptomic findings combined with our pharmacological data support a predominant role for β1-adrenergic receptor signaling in epinephrine-stimulated hormone release in human L-cells. These findings identify a previously underappreciated pathway through which circulating adrenergic signals may regulate GLP-1 and PYY secretion from the L-cells within the ileum of the human intestine.

The magnitude and spatial extent of the Ca²⁺ responses elicited by acetylcholine and epinephrine were substantially greater than expected from the abundance of L-cells in our organoids, raising the possibility that stimulation of a small population of sensor cells initiates a signal that propagates through the neighboring epithelium. Gap junctions provide a mechanism for direct intercellular propagation of ions and second messengers, and pharmacological disruption of intercellular coupling can suppress multicellular Ca²⁺ propagation in a range of epithelial and excitable tissues. We therefore tested whether the organoid-wide responses observed following cholinergic and adrenergic stimulation depended on intercellular coupling.

Pretreatment with the gap-junction inhibitor carbenoxolone progressively restricted the spatial extent and magnitude of the acetylcholine-induced Ca²⁺ response (**Fig. 5A,B**). At 100 μM carbenoxolone, the widespread epithelial wave was largely abolished, while discrete cells or small regions of the epithelium retained acute Ca²⁺ responses to acetylcholine. Despite this marked reduction in whole-organoid Ca²⁺ signaling, acetylcholine continued to stimulate GLP-1 secretion across all the concentrations of carbenoxolone tested (**Fig. 5C**). Moreover, epinephrine showed the same dissociation between epithelial Ca²⁺ propagation and endocrine output. Carbenoxolone reduced the magnitude and spatial extent of the epinephrine-induced Ca²⁺ response in a concentration-dependent manner, leaving spatially restricted responding cells at the highest concentration tested (**Fig. 5D,E**). Nevertheless, epinephrine continued to stimulate both GLP-1 and PYY secretion in the presence of carbenoxolone (**Fig. 5F**). These results indicate that the intercellular Ca²⁺ wave that propagates across the intestinal epithelium is not required for cholinergic or adrenergic stimulation of L-cell hormone release in human ileal organoids.

**Fig. 5.**
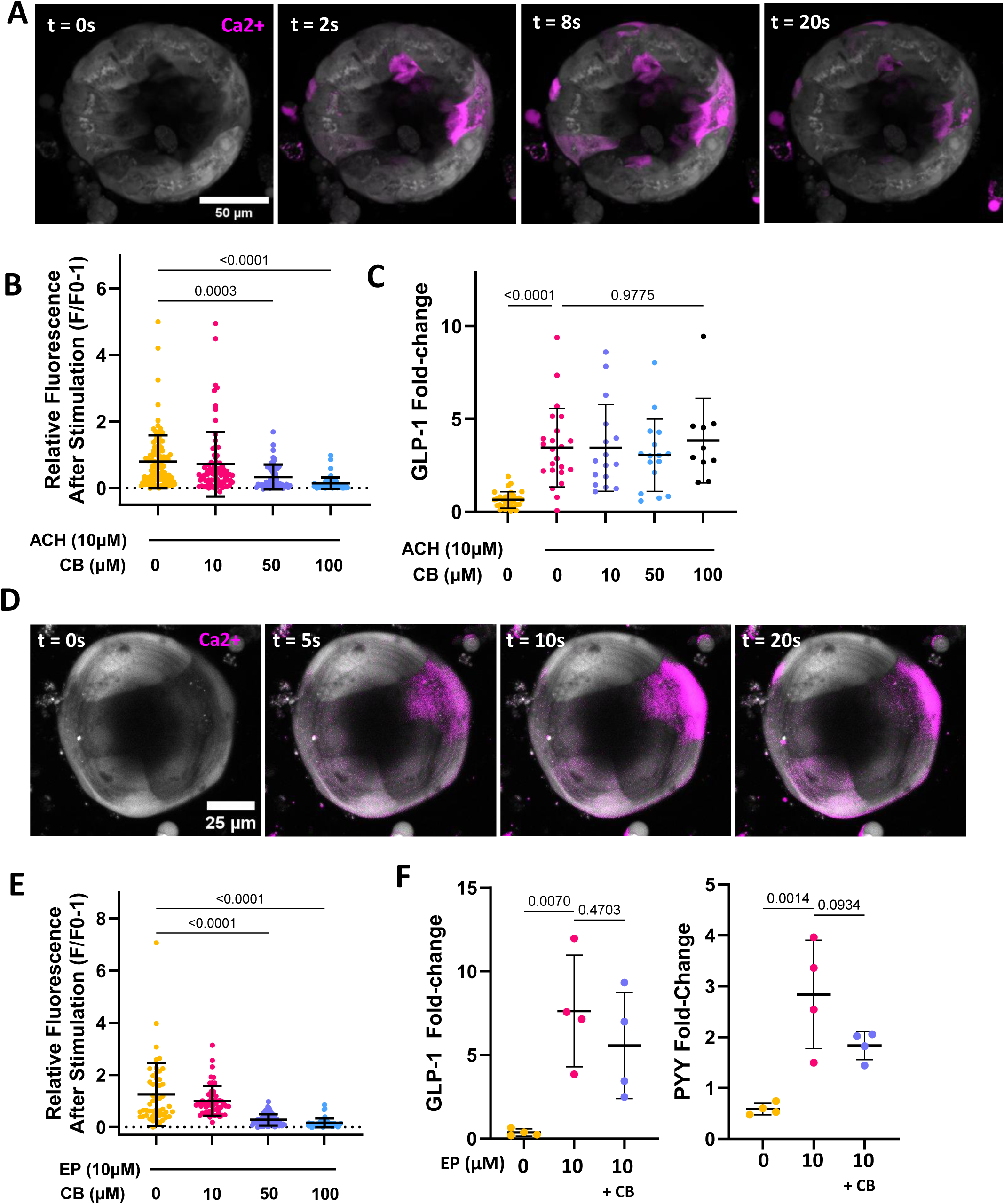
Epithelial calcium waves are dispensable for cholinergic and adrenergic stimulation of L-cell hormone secretion. **(A)** Representative maximum intensity projection time-lapse Ca²⁺ imaging of an organoid pretreated with 100 μM carbenoxolone (CB) and stimulated with 10 μM acetylcholine (ACH). t = 0 image shows baseline fluorescence immediately before stimulation; magenta indicates stimulus-evoked increase in Ca²⁺-dependent fluorescence at the indicated time points. Scale bar = 50 μm. **(B)** Peak Ca²⁺ response to 10 μM ACH following pretreatment with the indicated concentrations of CB. Increasing concentrations of CB progressively reduced whole-organoid Ca²⁺ responses (Lines indicate SD, comparison by one-way ANOVA with Dunnett’s multiple-comparison test versus ACh without CB, nonsignificant results not shown, n = 20–48 organoids from 3-5 independent wells per conditions across two organoid lines). **(C)** Within-well fold change in GLP-1 secretion following stimulation with 10 μM ACH in the presence of the indicated concentrations of CB. ACh significantly increased GLP-1 secretion, and secretion was not significantly reduced by CB over the concentration range tested (Lines indicate SD, comparison by one-way ANOVA with Dunnett’s multiple comparison test versus ACH without CB, n = 10–37 across two organoid lines). **(D)** Representative maximum intensity projection time-lapse Ca²⁺ imaging of an organoid pretreated with 100 μM CB and stimulated with 10 μM epinephrine (EP). t = 0 image shows baseline fluorescence immediately before EP stimulation; magenta indicates stimulus-evoked increase in Ca²⁺-dependent fluorescence at the indicated time points. Scale bar = 25 μm. **(E)** Peak Ca²⁺ response to 10 μM EP following pretreatment with the indicated concentrations of CB. Increasing concentrations of CB progressively reduced whole-organoid Ca²⁺ responses (Lines indicate SD, comparison by one-way ANOVA with Dunnett’s multiple-comparison test versus EP without CB, n = 50–59 organoids from 4 independent wells per condition). **(F)** Within-well fold change in GLP-1 and PYY secretion following stimulation with 10 μM EP in the absence or presence of 100 μM CB (Lines indicate SD, comparison by one-way ANOVA with Dunnett’s multiple-comparison test versus vehicle; n = 4).

Enteroendocrine hormone secretion is tightly coupled to cellular excitation and elevations in intracellular Ca²⁺. Primary L-cells are electrically excitable, and secretagogues that raise intracellular Ca²⁺ can drive GLP-1 release through Ca²⁺-dependent exocytotic machinery^18^. Furthermore, blocking the Ca²⁺ response is known to block hormone secretion in L-cells^17^. The persistence of GLP-1 and PYY secretion when epithelial Ca²⁺ propagation is suppressed therefore suggests that the residual stimulus-responsive cells include the hormone-secreting L-cell population.

Taken together with the muscarinic and β-adrenergic receptor pharmacology described above, these findings support a model in which acetylcholine and epinephrine act on a restricted epithelial sensor population that includes L-cells, triggering gut hormone secretion while secondarily initiating a broader wave of Ca²⁺ signaling through neighboring epithelial cells. Thus, cholinergic and adrenergic stimulation produce two separable outputs: an endocrine response that drives GLP-1 and PYY secretion, and a propagated epithelial Ca²⁺ wave whose physiological function remains to be determined.

## Discussion

We identified neural mechanisms that regulate human intestinal L-cell function and gut hormone secretion using a human donor-derived ileal organoid culture model that we optimized for enteroendocrine L-cell differentiation. Our studies revealed that cholinergic and adrenergic signals evoke epithelial calcium responses in human L-cells and stimulate secretion of two key metabolic hormones: GLP-1 and PYY. Neurotransmitter-evoked activation initiates carbenoxolone–dependent calcium waves that propagate through multiple cells across the intestinal epithelium; however, inhibition of calcium wave propagation does not impair hormone release. These findings suggest that L-cells are key cellular targets of these neurotransmitters and that the associated epithelial calcium waves reflect coordinated downstream responses rather than a requirement for endocrine hormone secretion.

Establishment of a human ileal organoid model with physiological levels of L-cells was critical in enabling this finding. Prior studies of neurotransmitter regulation of L-cells have relied on rodent models or immortalized neuroendocrine cell lines, systems in which receptor expression and signaling properties may differ substantially from those of human intestinal epithelium. Typical models of the human ileum have extremely low levels (< 1%) of L-cells, making measurement of L-cell hormone secretion challenging without the addition of cAMP activators like forskolin and IBMX^39,42–44^. Other groups have attempted to increase L-cell numbers through genetic engineering with mixed results.^45,46^ Notably, by modifying enteroendocrine differentiation conditions established for rectal organoids through additional inhibition of EZH2 using EPZ, we achieved a multi-fold increase in L-cell yield compared with existing human organoid protocols. This improvement allowed ligand-evoked secretion to be quantified without prior pharmacologic elevation of cAMP, avoiding a pre-stimulated secretory background. This enrichment also allowed direct measurement of calcium dynamics and hormone secretion in donor-derived ileal organoids, which provided a platform for defining human-specific regulatory mechanisms governing gut hormone release. This model also offers promise to better understand human disease linked to L-cell function, such as diabetes and obesity.

Importantly, our findings establish a neural pathway by which cholinergic signals regulate human L-cell activity. Our pharmacological analyses demonstrate that acetylcholine-induced calcium responses and hormone secretion profiles are consistent with a predominant role for M3 muscarinic receptor signaling, with little contribution from nicotinic signaling. These findings contrast with prior observations in rodent systems and cell lines, where both receptor classes have been implicated in GLP-1 release, and they highlight the importance of studying neurotransmitter regulation in primary human L-cell models^28,47^. Given the dense cholinergic innervation of the distal small intestine^48,49^, these findings suggest that muscarinic signaling may represent a key route through which enteric and parasympathetic activity during feeding and digestion directly modulate L-cell hormone output to coordinate nutrient absorption with endocrine control of metabolism^9,50^.

Interestingly, in addition to cholinergic signaling, we found that epinephrine activated human L-cells to induce calcium influx and hormone secretion through the β1-adrenergic receptor. Unlike acetylcholine, epinephrine is unlikely to be released locally by enteric neurons, suggesting that L-cell responsiveness to epinephrine may reflect a sensitivity to circulating catecholamines. In this context, L-cells may function as sensors of systemic autonomic state, coupling stress- or arousal-associated adrenergic signals to intestinal hormone secretion. Thus, the convergence of cholinergic and adrenergic pathways at the level of the L-cell provides a mechanism by which gut hormone output can be dynamically tuned to both local digestive cues and organism-wide metabolic demands.

The identification of β1 adrenergic receptor signaling in cultured human ileal organoids also raises the possibility that norepinephrine contributes to L-cell regulation *in vivo*. Norepinephrine is the principal catecholamine released by postganglionic sympathetic neurons^51^, and prior work has identified close anatomical apposition between sympathetic fibers and intestinal L-cells^21^. In addition, sympathetic adrenergic signaling has been shown to regulate L-cell calcium activity and GLP-1 secretion through α2A receptors in mice, establishing the existence of a functional sympathetic nerve–L-cell axis at least in rodents^21^. Norepinephrine and epinephrine both activate β1-adrenoceptors and structural studies indicate that norepinephrine exhibits substantial selectivity for β1 over β2 receptors^52^. Although norepinephrine did not produce a statistically significant response in our whole organoid assays, bath application may not reproduce the spatially restricted catecholamine exposure generated by sympathetic nerve terminals *in vivo*. Our findings therefore do not establish a physiological role for norepinephrine, but raise the possibility that local sympathetic signaling could engage β1-dependent stimulatory pathways under conditions not captured by the organoid culture model used in this study.

Our finding that β1 receptor activation promotes GLP-1 and PYY secretion by human L-cells differs from studies of perfused rat ileum that implicated β2 receptors as the predominant stimulatory β-adrenergic subtype^53^, as well as from recent work identifying α2A-mediated sympathetic inhibition of L-cell activity *in vivo*^21^. However, β1 receptor-dependent stimulation of GLP-1 exocytosis has also been reported in GLUTag cells, where α1, β1, and α2 receptor pathways exerted opposing effects on secretion^54^. Together, these findings suggest that adrenergic control of L-cell function may not be uniformly stimulatory or inhibitory, but instead depend on the relative expression and engagement of adrenergic receptor subtypes. Differences among species, intestinal regions, experimental preparations, catecholamine concentrations, and physiological context may therefore shape the net effect of sympathetic signaling on GLP-1 release.

An unexpected finding of this study is that neurotransmitter-induced activation initiates calcium waves that propagate through the surrounding intestinal epithelium. Pharmacological inhibition of gap junctions suppressed calcium wave propagation without altering calcium responses with the subset of epithelial cells that initiate the response, suggesting that calcium signals propagate between cells via an intracellular route. Hormone secretion from L-cells is coupled to increases in intracellular calcium^38^, but interference with propagation of the calcium wave using a gap junction inhibitor did not interfere with hormone secretion. These results indicate that the epithelial calcium waves that are triggered by the neurotransmitters we observed are not required for endocrine output. This finding that hormone secretion continues during treatment with carbenoxolone demonstrates that L-cells are directly activated by acetylcholine and epinephrine, rather than downstream targets of the propagating calcium wave. The functional significance of these calcium waves remains to be determined; however, one possibility is that they coordinate intestinal epithelial behavior in response to neural input, analogous to calcium waves described in other epithelial systems that regulate processes such as secretion, barrier function, or tissue repair^55–60^. Identifying the specific connexin isoforms involved and defining the downstream consequences of epithelial calcium propagation will be important areas for future investigation.

Our data also raise the possibility that cholinergic regulation of L-cells is not exclusively neuronal in origin. In addition to enteric neurons, other epithelial cell types (e.g., tuft cells) have been shown to produce acetylcholine in response to microbial and inflammatory cues^61^. This suggests that cholinergic activation of L-cells may integrate neuronal, epithelial, and immune-derived signals within the intestinal mucosa, further reinforcing the role of L-cells as central integrators of local and systemic information relevant to metabolic homeostasis.

Several limitations of this study should be noted. The organoid culture system lacks intact neural circuitry, vascular inputs, and mechanical forces associated with peristalsis that are present in vivo, which likely further increase the spatial and temporal complexity of intestinal signaling. Moreover, the receptor subtype assignments are based on pharmacological selectivity rather than cell-specific genetic perturbation, and the identity of the residual calcium-responsive cells after inhibition of intercellular propagation remains to be directly established. Finally, while GLP-1 and PYY secretion were robustly induced by neural and neuroendocrine stimuli, additional L-cell–derived peptides could not be reliably quantified, leaving open questions regarding the broader hormonal output elicited by these pathways.

In summary, our findings revealed cholinergic and adrenergic mechanisms that regulate human intestinal L-cell activity and gut hormone secretion. By positioning L-cells as integrative hubs that translate autonomic and neuroendocrine signals into coordinated epithelial and endocrine responses, this work provides mechanistic insight into how nervous system state is coupled to intestinal hormone release in humans. These results have implications for understanding communication through the gut–brain axis and for development of alternative therapeutic strategies aimed at modulating GLP-1 and related hormones to treat metabolic disease.

## Methods

### Human tissues and organoid lines

Human intestinal organoid lines were generated from de-identified biopsies obtained from patients undergoing endoscopy at Boston Children’s Hospital or surgical resections at Massachusetts General Hospital. Samples were taken from tissue regions without gross pathology. Consent from guardians and assent from donors were collected in accordance with IRB approval (Boston Children’s hospital protocol number IRB-P00000529, Massachusetts General Hospital protocol number IRB-2015P001859). Organoid lines were derived from donors aged 11-59 years (n=4; 2 male, 2 female). Donor age and sex were unknown to researchers when experiments were being performed.

Biopsies were either processed immediately in DMEM/F12 or cryopreserved in 10% FBS with DMSO. To generate organoids, intestinal crypts were isolated from fresh or thawed samples by rinsing in DMEM/F12 and digesting in 2 mg/mL Collagenase Type I prepared in Hank’s Balanced Salt Solution.

Tissue fragments were agitated with a motorized pestle, then incubated at 37 °C for 40 minutes. At the midpoint, digestion was aided by pipetting with a bent pipette tip. Following incubation, samples were centrifuged at 800 × g for 5 minutes and supernatants were aspirated. Crypts were resuspended in 150 μL Matrigel, with 40 μL seeded into three wells of a 24-well plate and polymerized at 37 °C for 10 minutes before culture medium was added.

### Organoid culture

Ileum organoids were grown in 45 µL matrigel (Corning) domes incubated at 37°C and 5% CO_2_ with 1 mL of growth medium (Supplementary Table 1) per well of a 24-well plate. Medium was changed every 2 – 3 days. Organoids were passaged weekly via mechanical dissociation. To do so, medium was aspirated and 500µL of ice cold cell recovery solution was added to each well. Matrigel domes were scraped, collected in cell recovery solution, and pooled in 15 mL tubes. Tubes were placed on ice and turned over every 10 to 15 minutes for 1 hour. Organoids were then spun down at 300g at 4 °C for 5 minutes, and the supernatant was aspirated. Organoids were resuspended in matrigel and dissociated by pipetting up and down 70 – 100 times with 1 mL pipette whose tip had been bent. Organoid fragments were then dispensed in 45 µL domes at the center of 24-well plates and polymerized in a 37 °C incubator for 15 – 25 minutes. After polymerization, 500 µL of growth medium with 10 µM Y-27632 rock inhibitor (Tocris) was added to each well. 24 hours later media was changed to 1 mL of growth media.

For differentiation, the same total number of organoids was resuspended in half the quantity of matrigel, and 25 µL matrigel domes were plated in 48 well plates. 500 µL of growth medium with 10uM Y-27632 was added to each well, and replaced after 24 hours with 500 µL growth medium. 48 hours after initial plating, media was changed to 500 µL differentiation medium (Supplementary Table 1), and differentiation medium was changed every two days. After twelve days differentiation was complete, except where otherwise noted. The L-WRN cell line was used to produce conditioned medium containing Wnt3a, Noggin and R-spondin 3, which was included in both the growth and differentiation media (Supplementary Table 1).

### RT-qPCR

Fully differentiated ileum organoids were extracted from matrigel using cell recovery solution, following the same procedure used for passaging. After spinning down and aspirating the supernatant, organoids were lysed in TRI reagent through repeated pipetting. RNA was extracted from lysates using a Zymo direct-zol RNA extraction kit (Zymo Research). RNA was converted to cDNA with a superscript IV reverse transcription kit (Thermofisher), and gene expression was measured by running qPCR with Taqman fast advanced master mix and Taqman primers (Supplementary Table 2) on a CFX96 RT-PCR detection system (Bio-Rad).

### Confocal Imaging

Organoids were extracted from matrigel following the same procedure used for passaging, resuspended in growth media with 10 µM Y-27632, and plated on Ibidi 96-well plates that had been coated with 2% matrigel for 1 hour at 37 °C and 5% CO_2_. Organoids were placed at 37 °C and 5% CO_2_ overnight to attach. The following day, organoids were fixed with a solution of 4% PFA in PBS +/+ for 20 minutes and washed 2x with PBS -/-. Cells were permeabilized with 0.25% Triton-X in PBS -/- for 5 minutes, and blocked with 5% donkey serum, 1% BSA and 0.25% Triton-X in PBS -/- for 1 hour.

Organoids were washed once with 0.01% Tween in PBS -/- (tween) and incubated overnight at 4oC in primary antibodies in tween (see Supplementary Table 3). The following day, organoids were washed 4x for 5 minutes in tween, and incubated at room temperature for two hours with secondary antibodies (Supplementary Table 3). Organoids were incubated in DAPI in tween for 10 minutes, then washed 2x with tween and 2x in PBS -/- before imaging in PBS -/-.

Images were collected on either a Leica Stellaris 8 using a 40X water immersion objective in the Leica Application Service X software or a Nikon Ti7 using a 20X air objective in the NIS Elements AR software. Images were analyzed in FIJI.

### Calcium imaging

Organoids were attached to Ibidi-96 well plates following the same procedure used for fixed imaging. After attaching overnight, organoids were washed once with HBSS +/+, and incubated with 5 µM Calbryte 520 AM calcium dye in HBSS +/+ at 37 °C and 5% CO_2_ for 45 minutes. Next, organoids were washed once with HBSS +/+, and imaged in 100 µL of HBSS +/+ using a Zeiss Axio 3 observer running in burst mode at 10Hz. For each trial, 10 seconds after beginning image collect, 100 µL of HBSS plus stimulus at double strength was pipetted into the well (see Supplementary Table 2 for stimulus). Total imaging time was 50 seconds. Positive controls were collected at the beginning and end of imaging to ensure organoids were viable throughout the imaging session.

Reported values for relative fluorescence after stimulation were measured on an organoid-by-organoid basis by drawing a region of interest around every organoid present in a frame, averaging fluorescence intensity across the frame for each time point, and subtracting an intensity of a background frame at the same time point to account for photobleaching of the media. For each organoid, the average of the first 5 seconds of data collected served as a baseline for fluorescence intensity. The rest of the time series was normalized to this baseline, and relative fluorescence after stimulation was calculated as the difference from the end of this 5 second period to the peak of fluorescence.

### Hormone secretion

Supernatants for hormone secretion measurements were collected from organoids grown and differentiated in matrigel droplets in 48 well plates. Each measurement consisted of a baseline supernatant and a study supernatant drawn sequentially from the same well. To generate the baseline supernatant, differentiation medium was removed from the well, the well was washed once with PBS -/-, aspirated dry, and 150 µL of basic medium was added. Basic medium consisted of advanced DMEM/F12 supplemented with 2% B-27, 1% N-2, 1% Glutamax, 1% HEPES (all sourced from Gibco), 500uM N-Acetyl-L-Cysteine (Sigma), and 100uM diprotin A (Tocris). Organoids were placed in the incubator for 3 hours, after which conditioned medium was collected to serve as a baseline supernatant. Wells were washed once with PBS -/-, aspirated dry, and 150 µL of basic medium supplemented with the appropriate stimulus or vehicle was added to the well (see Supplementary Table 2 for stimulus). Organoids were returned to the incubator for 3 hours, and conditioned medium was collected to serve as study supernatant. In conditions where inhibitors or antagonists were used, an additional 30 minute incubation with basic medium supplemented with inhibitors or vehicle was added between baseline and study collections to ensure that inhibitors were present before the addition of any stimulus.

Resulting supernatants were frozen at -80 °C. Measurement of GLP-1 and PYY was carried out using Mesoscale Discovery U-Plex kits coated with detection antibodies for human GLP-1 (total) and human PYY. Kits were run according to manufacturer instructions, and analyzed using a Meso Quickplex SQ 120. Reported fold-change values for GLP-1 and PYY were generated by dividing the calculated concentration of the study supernatant by the baseline supernatant on a well-by-well basis.

### Gene expression

Raw counts were obtained from GEO (GSE284419, GSE114853, GSE148224, GSE146799) and analyzed in R 4.4.3 with DESeq2^62^. Ensembl identifiers were mapped to gene symbols and counts for duplicate symbols summed. Genes with zero counts across all samples of a dataset were removed. Counts were normalized by the DESeq2 median-of-ratios size factors. Sorted populations were harmonized to L-cell, other EEC and non-EEC. Fold changes were shrunken with apeglm^63^. p-values represent two-sided Wald tests on the unshrunken estimates with Benjamini-Hochberg adjustment across all genes; adjusted p < 0.05 was considered significant. Genes removed by DESeq2 independent filtering receive no adjusted p-value and are reported as untested. Figures were produced with ComplexHeatmap^64^. All data are public and available from GEO under the accessions above.

Models follow each study’s design. GSE114853: ∼ participant + tissue + population, as all 11 donors contributed all three sorted populations. GSE148224: ∼ prep + population, as each organoid preparation contributed matched Venus+ and Venus− samples, with preparation nested within growth medium. GSE146799: ∼ population. All samples of a dataset were retained in every fit so that dispersion is estimated across the full design; the reference level was reset and the model refitted for each contrast.

GSE284419 is a single-cell dataset and comprises one donor. Cells were processed in Seurat^65^, retaining those with 500–9,000 detected genes and <25% mitochondrial reads, then log-normalized, reduced over 2,000 variable features and 30 principal components, and clustered at resolution 1.0. Clusters were labelled from mean marker expression: non-EEC where the mean of CHGA, CHGB, SCG2, SCG3 and PCSK1N fell below 1.0, and L-cell where mean GCG exceeded 1.0 (a clean gap separated cluster means of 0.68 and 1.60). Fold changes are pseudo-bulk CPM ratios, log2((CPM_L-cell + 1)/(CPM_comparator + 1)). p-values were not computed.

## ACKNOWLEDGEMENTS

The authors would like to thank Girija Goyal, Kenneth Carlson, and Sylvie Bernier for their help developing the experimental approach and interpretation of the results, and Tom Ferrante for his help setting up and troubleshooting imaging systems.

## AUTHOR CONTRIBUTIONS

J. L. designed and performed experiments, analyzed data, prepared figures, and wrote the manuscript; D.Z. provided conceptual advice, generated organoid lines, performed experiments, and revised the manuscript; S.R. performed experiments and revised the manuscript; A. S. provided conceptual advice, interpreted data, and revised the manuscript; A. G. performed computational analysis and revised the manuscript; D. B. designed experiments, provided conceptual advice, and provided organoid lines; M. L. supervised the work, interpreted data, and provided conceptual advice; D. I. supervised the work, designed experiments, interpreted data, revised the manuscript, and acquired funding.

## COMPETEING INTERESTS

The authors declare no competing interests for this work.

## DATA AVAILABILITY

All data generated in support of the findings of this study are available within the paper. Raw data are available upon request. External datasets referenced can be found at the references linked in the text.

