## Supplemental Tables for "Neural Activation of Human Primary Intestinal Enteroendocrine L-cells"

| Supplementary Table 1 - Media Formulations | |
| --- | --- |
| Growth Medium | Differentiation Medium |
| L-WRN conditioned medium (65% v/v) | |
| Advanced DMEM/F12 (30% v/v/) | |
| Glutamax (1% v/v) | |
| B-27 (1% v/v) | |
| N-2 (0.5% v/v) | |
| HEPES (10 mM) | |
| Primocin (100 μg/mL) | |
| Normocin (100 μg/mL) | |
| A83-01 (500 nM) | |
| N-acetyl-L-cystein (500 μM) | |
| murine EGF (50 ng/mL) | |
| Human [Lue] Gastrin I (50 nM) | |
| Nicotinamide (10 mM) | DAPT (20 μM) |
| SB202190 (10 μM) | Betacellulin (20 ng/mL) |
|  | PF06260933 (6 μM) |
|  | Tranylcypromine (1.5 μM) |
|  | EPZ 6438 (1 μM) |
|  | Prostaglandin E2 (10 nM) |

**Supplementary Table 1.**

Growth and Differentiation Media Components Growth factors, supplements, and small molecules common to both growth and differentiation medias are listed first. The L-WRN cell line was used to produce conditioned media containing Wnt3a, Noggin and R-spondin 3.

| Supplementary Table 2 - Compound list | |  |
| --- | --- | --- |
| Compound | Supplier | Catalog Number |
| Acetylcholine Chloride | Selleckchem | S1805 |
| Norepinephrine bitartate | Tocris | 5169 |
| Epinephrine HCl | Selleckchem | S3061 |
| GABA | Selleckchem | S4700 |
| Vasoactive Intestinal Peptide | Echelon biosciences | 491-25 |
| Bethanechol Chloride | Selleckchem | S2455 |
| (−)-Nicotine hemisulfate salt | Sigma Aldrich | N1019 |
| Atropine sulfate monohydrate | Selleckchem | S2130 |
| Pirenzepine dihydrochloride | Selleckchem | S3662 |
| Methoctramine hydrate | Sigma Aldrich | M105-10MG |
| Darifenacin | Selleckchem | E7262 |
| PD 102807 | Tocris | 1671 |
| Isoproterenol | Selleckchem | S2566 |
| ICI 118,551 | Selleckchem | S8114 |
| CGP 20712 dihydrochloride | Tocris | 1024 |
| Terbutaline hemisulfate | Selleckchem | S4127 |
| Phenylephrine | Selleckchem | S2569 |
| Carbenoxolone disodium | Sigma Aldrich | C4790-1G |

| Supplementary table 3 - Antibodies and Stains | |  |
| --- | --- | --- |
| Compound | Supplier | Catalog Number |
| CHGA | Cell Signaling Technology | 85798 |
| GLP-1 | abcam | ab23468 |
| LYZ | Novus Biologicals | NB100-63062 |
| MUC2 | Novus Biologicals | NBP2-44430-0.1mg |
| 5-HT | abcam | ab66047 |
| Ki67 | abcam | ab15580 |
| Laminin | Novus Biologicals | NB300-144 |
| Donkey Anti-Rabbit Alexa Fluor Plus 555 | Invitrogen | A32794 |
| Donkey Anti-Goat Alexa Fluor Plus 594 | Invitrogen | A32758 |
| Donkey Anti-Mouse Alexa Fluor Plus 647 | Invitrogen | A32787 |
| DAPI | Invitrogen | D1306 |
| Phalloidin | Invitrogen | A12381 |
